# Automated methods for 3D Segmentation of Focused Ion Beam-Scanning Electron Microscopic Images

**DOI:** 10.1101/509232

**Authors:** Brian Caffrey, Alexander V. Maltsev, Marta Gonzalez-Freire, Lisa M. Hartnell, Luigi Ferrucci, Sriram Subramaniam

## Abstract

Focused Ion Beam Scanning Electron Microscopy (FIB-SEM) is an imaging approach that enables analysis of the 3D architecture of cells and tissues at resolutions that are 1-2 orders of magnitude higher than that possible with light microscopy. The slow speeds of data collection and analysis are two critical problems that limit more extensive use of FIB-SEM technology. Here, we present a robust method that enables rapid, large-scale acquisition of data from tissue specimens, combined with an approach for automated data segmentation using machine learning, which dramatically increases the speed of image analysis. We demonstrate the feasibility of these methods through the 3D analysis of human muscle tissue by showing that our process results in an improvement in speed of up to three orders of magnitude as compared to manual approaches for data segmentation. All programs and scripts we use are open source and are immediately available for use by others.

**Impact Statement:** The high-throughput, easy-to-use and versatile segmentation pipeline described in our manuscript will enable rapid, large-scale statistical analysis of sub-cellular structures in tissues.

## INTRODUCTION

Focused Ion Beam Scanning Electron Microscopy (FIB-SEM) is an approach for 3D imaging of specimens with thicknesses greater than ~ 1 micron that cannot be imaged using transmission electron microscopy due to their thickness. In biological FIB-SEM imaging, a focused gallium ion beam is used to progressively remove material from the surface of a macroscopic specimen such as a cell pellet or tissue specimen, with the recording of a backscattered electron microscopic image using a scanning electron beam. The resulting volumes contain useful information on subcellular architecture at spatial resolutions as high as ~ 10 nm, and visualization of the data in 3D by segmentation can provide new and unexpected insights into the organization of organelles and membranes in the cell. (Glancy, Hartnell and Combs, et al. 2017; Glancy, Hartnell and Malide, et al. 2015; Narayan and Subramaniam 2015).

As currently used, the speed of interpreting the image stack using segmentation approaches to delineate membrane and organelle boundaries is the principal bottleneck in the application of FIB-SEM. To realistically address biologically and medically interesting problems, increases in the speed of segmentation of at least two orders of magnitude are required. A recent estimate put the amount of time required, using present approaches for manual segmentation, to segment a 1×10^5^ um^3^ volume to take between 2×10^4^ - 1×10^5^ work hours to complete (Berning, Boergens and Helmstaedter 2015), this is not including the time taken to acquire such large volumes at high resolutions in the first place.

Machine learning, and other advanced computational techniques have begun to dramatically reduce the time taken to convert an imaged volume to discrete and quantifiable structures within the volume (Januszewski, et al. 2018; Berning, Boergens and Helmstaedter 2015; Lucchi, et al. 2011; Meijs, et al. 2017; Camacho, et al. 2018; Kasaragod, et al. 2018). The approach we present here takes an integrative, high-throughput and easy-to-use approach towards sample collection, segmentation and analysis with the view to creating a versatile but accurate methodology for tackling a multitude of biologically relevant problems.

The class of problems we are interested in requires comparison of the 3D distribution of mitochondria in muscle tissues obtained from human volunteers of different ages, with the goal of combining this information with biochemical and proteomic analyses of the samples to define the biology of aging. We estimate that to obtain a statistically meaningful analysis of human tissues we would require several volumes of an individual’s muscle fibers (see Figure 1), each >1×10^4^ um^3^ in size across many individuals dispersed over a wide age range. Using this approach, to discern potential age-related variations in mitochondrial architecture with manual segmentation could take years. The data collection approaches used here enabled collection from 3.44×10^5^ μm^3^ of human skeletal muscle samples from four healthy male individuals in ~12 instrument days. Segmentation of the data sets was achieved at the rate of −2800 μm^3^ per hour, corresponding to a ~500-3000-fold increase in speed relative to manual segmentation without loss of the detail required to interpret the image data.

**Figure 1.**
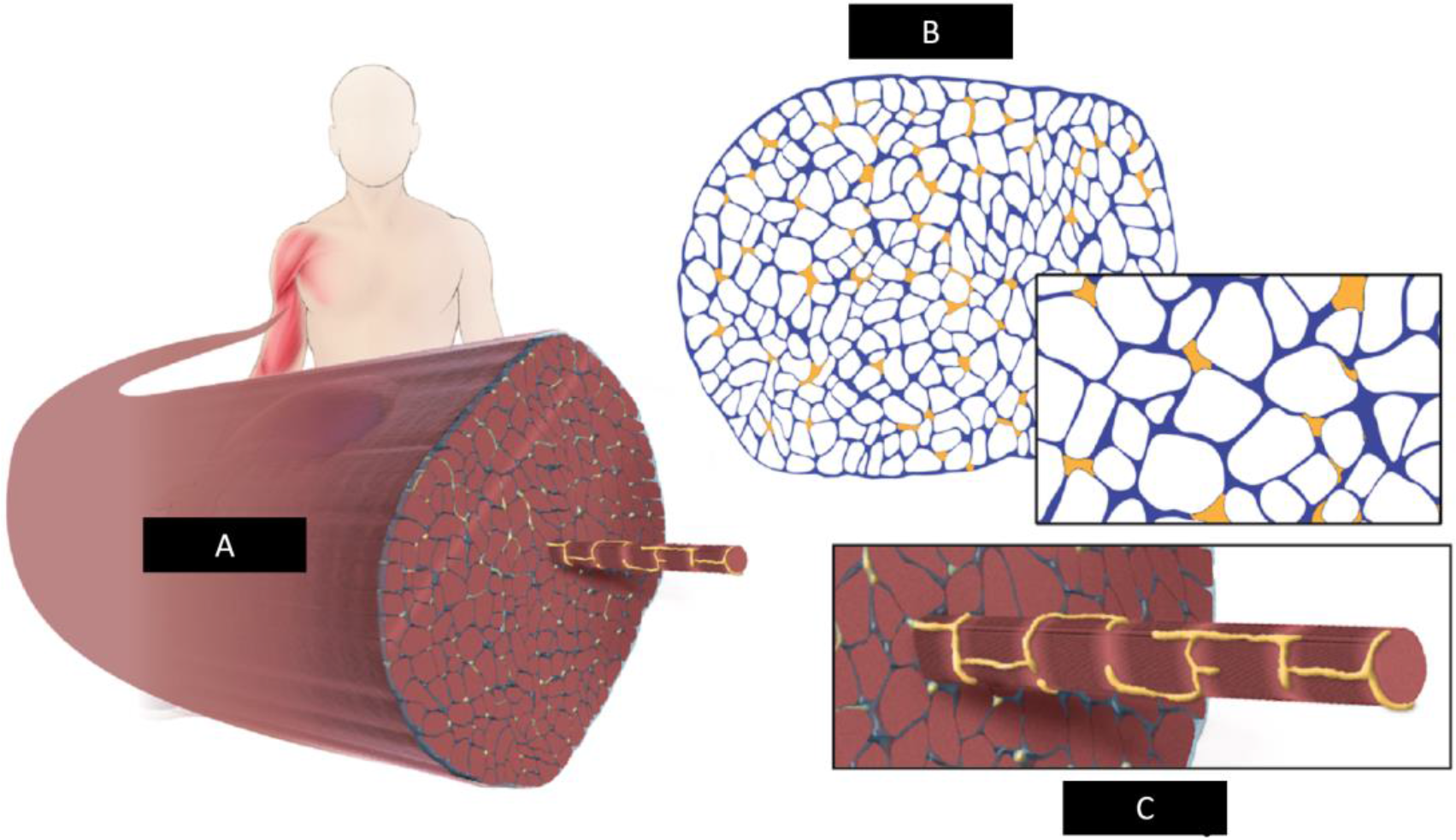
Illustration of human skeletal muscle fiber. **A)** Whole Skeletal Muscle Fiber. **B)** Muscle Fiber cross-section. **C)** Individual myofibril.

## RESULTS

### Workflow for data collection and segmentation

In Figure 2, we present orthogonal views of the SEM images from a representative FIB-SEM data collection run with a muscle tissue specimen. Early in the design of our experiments, we found that increasing the voxel size from 5×5×15nm^3^ to 15×15×15nm^3^ increased the rate of data acquisition ~3-fold from ~500 μm^3^ per hour to ~1500μm^3^ per hour. We established that the information required for segmentation was not compromised by the use of larger pixel sizes in the x and y dimensions (Figure 2 - Supplemental Figure 1) for the purpose of recognizing mitochondria.

**Figure 2.**
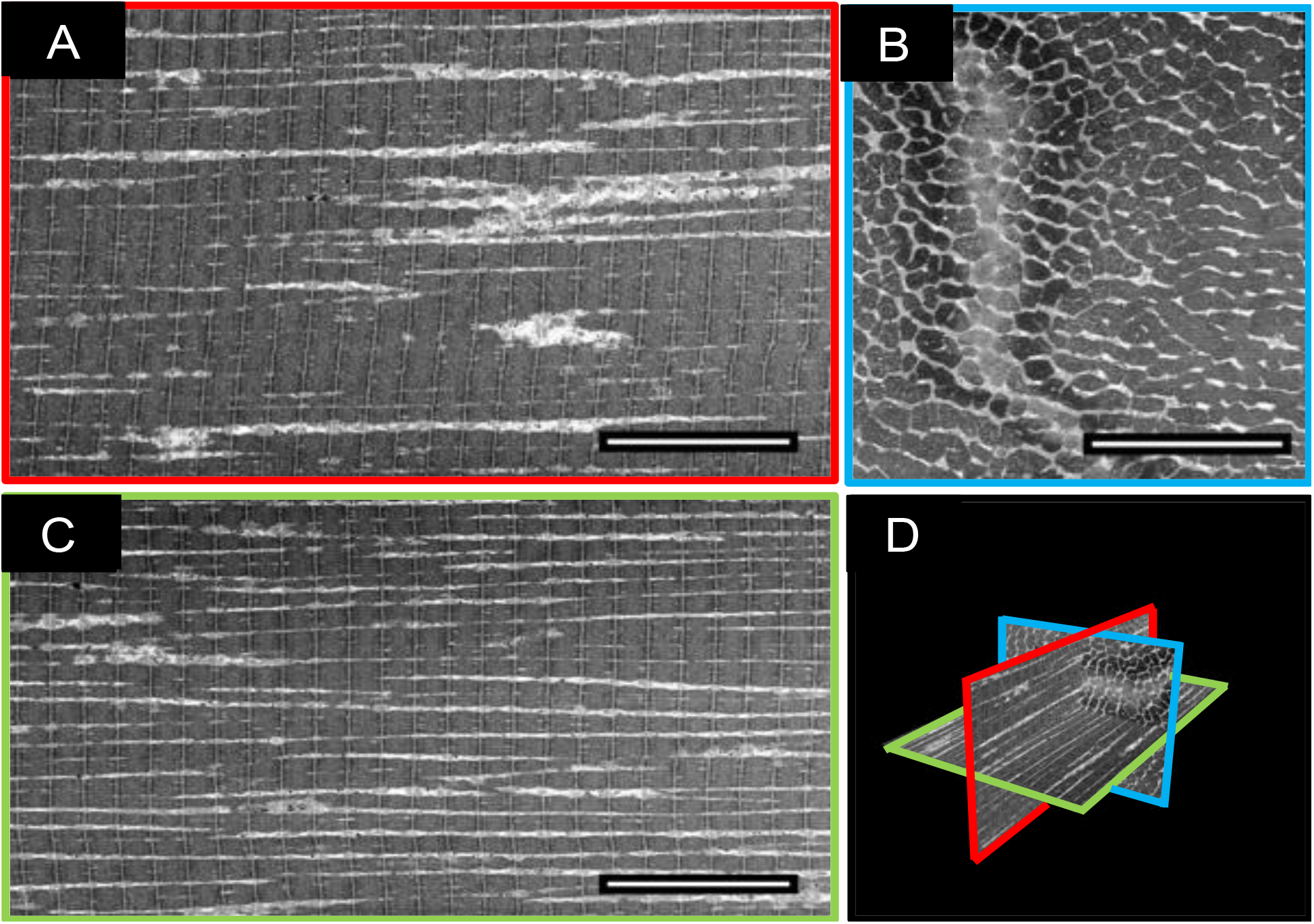
A typical example of a muscle fiber acquired with FIB-SEM at a voxel size of 15nm^3^. **A)** Z-Axis (Imaging) face of FIB-SEM volume. **B)** X-Axis face of FIB-SEM volume. **C)** Y-Axis face of FIB-SEM volume. **D)** 3D Orthoslice representation of slices A-C. Scale Bar = 10μm.

Manual analysis of the 3D image stack shows that the mitochondria display two distinct architectural arrangements, with one class displaying thick, densely packed networks (type A) and those with thin, sparse networks (type B). The overall spatial arrangements of these mitochondrial types (Figure 3) are distinct, with the type A fibers (Figures 3A, 3B) forming a highly connected assembly, while the type B fibers (Figures 3C, 3D) are arranged in smaller clusters in addition to being loosely packed. See Figure 3 - Supplemental Figure 1, 2 for video of Type A and B 3D segmentation respectively.

**Figure 3.**
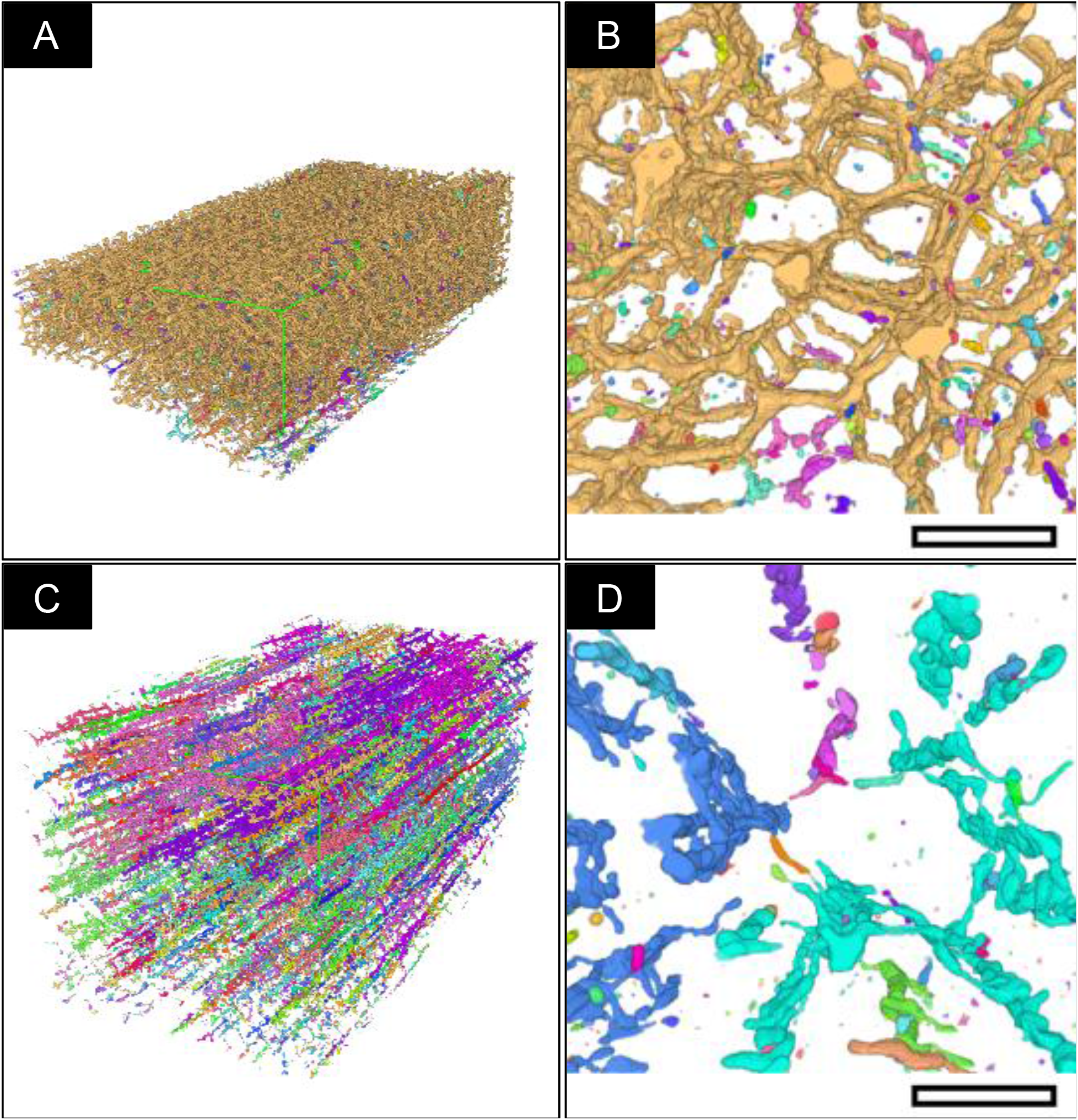
Morphological Classification. Each continuous network of connected mitochondria, as determined by ImageJ’s “MorphoLibJ” plugin, in the above images were labelled a single color. **A)** Typical “Type A” fiber segmentation volume. **B)** Transverse “Type-A” (X-Axis) image of a mitochondrial sub-volume. The majority of mitochondria in this volume are from a single network, indicated by a uniform label across the whole volume. **C)** Typical “Type B” fiber segmentation volume **D)** Transverse “Type B” (X-Axis) image of mitochondrial sub-volume. The majority of mitochondria in this volume are from multiple discontinuous networks indicated by the multi-colored labelling evident in the volume. Scale bar: 2 μm.

We combined volume acquisition, alignment and normalization, machine learning (ML) training, automated segmentation and statistical analysis into a pipeline and used it to segment multiple tissue volumes (Figure 4).

**Figure.4.**
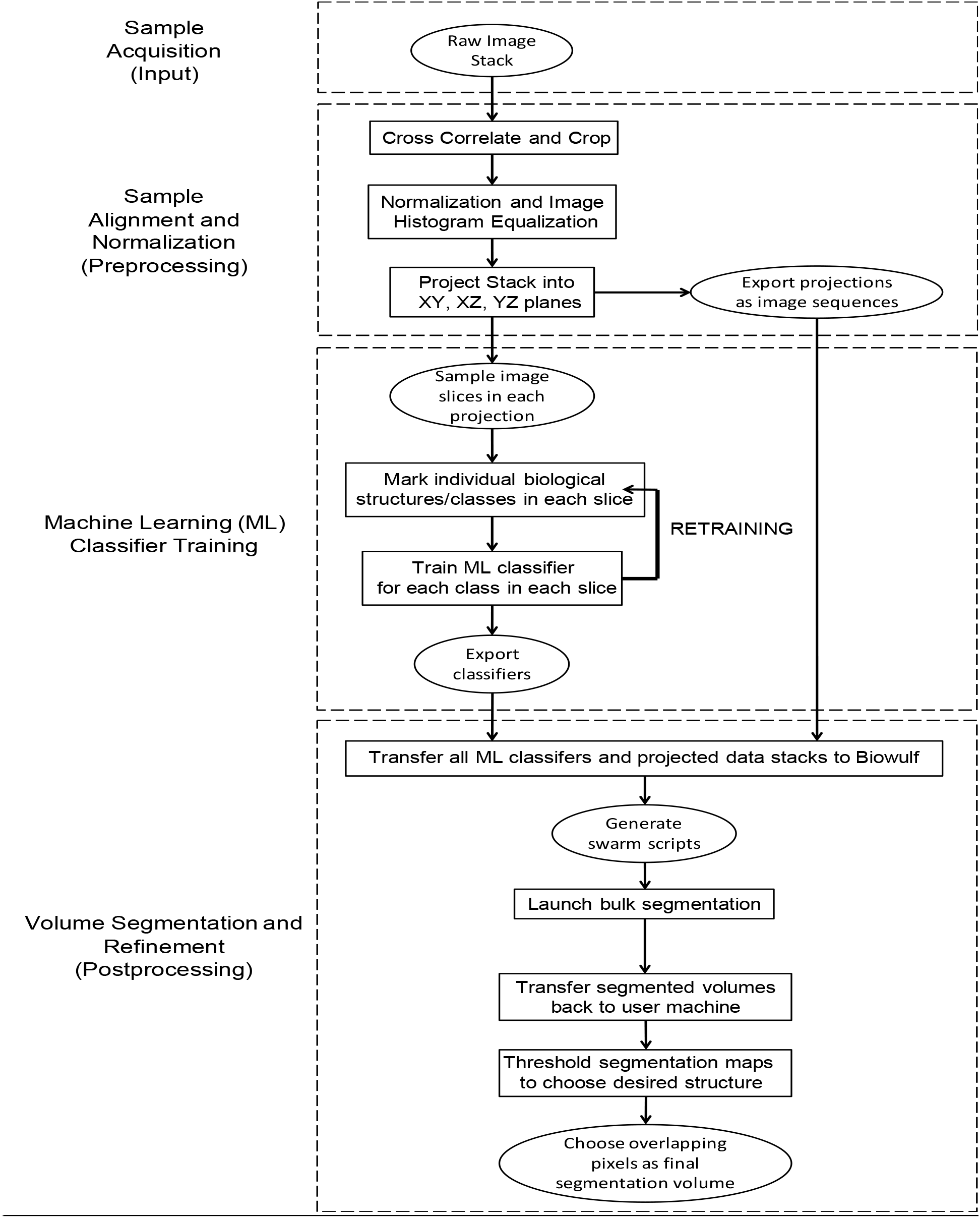
Segmentation Pipeline. Flowchart indicating the significant steps in the acquisition, segmentation and analysis of 3D volumes.

Below, we summarize the main steps of our approach:

1. <u>Sample Acquisition (Input):</u> Once a suitable area was found on a given tissue block, it was imaged with a 15×15×15nm voxel size (xyz) resulting in final volume dimensions of 60×30×30 μm^3^ (54,000μm^3^).
2. <u>Sample Alignment and Normalization (Preprocessing):</u> The individual images (in tiff format) were aligned to a complete 3D stack using a cross-correlation algorithm as described previously (Murphy, et al. 2011). The resulting (.mrc) file was opened in ImageJ cropped, median filtered, binned to a voxel size of 30nm^3^ and the stack histogram was normalized and equalized for reproducibility between volumes. 3 slices were selected from each of the principal axes, evenly spaced across the volume, resulting in 9 images for manual classification.
3. <u>Machine Learning Training (Manual Classification):</u> Each of the major biological structures in the 9 images were classified based on their standard histological features (z-disk, mitochondria, A-band, I-band, sarcoplasmic reticulum and lipids). After sufficient annotation, the Weka segmentation platform was used to train the machine learning software on the images. The output was inspected, and if the software failed to classify the image adequately, the above classification process was repeated iteratively until the software produced an accurate classification of the slice. A (.model) file was then exported to the biowulf computing resource at the NIH (details on how to port the Weka segmentation platform to a generic computing cluster are included in the supplementary information).
4. <u>Volume Segmentation and Refinement (Postprocessing):</u> The volume prepared in step 2 was exported to biowulf as a series of individual image slices, and each of the 9 classifiers were applied to the image stack, producing 9 x 32-bit tiff format outputs of classified images, which were then imported from biowulf and processed on local computers using the ImageJ image processing package. Images classified as mitochondria were isolated as binary 8-bit tiff format files. The 3 volumes from each axis were first added together using ImageJ’s “Image calculator” function; densities that did not overlap with at least one of the other 2 volumes were removed through simple thresholding. Each axis volume was then added together using the previously mentioned function, and density which did not overlap with at least one of the other 2 axes was removed through simple thresholding. The resulting 3D volume, after low-pass filtering was used for statistical analysis of mitochondrial densities.
5. <u>Statistical Analysis:</u> Each volume was classified according to the mitochondrial density pattern into either a “type A” or “type B” fiber (as defined in Figure 3). The resulting average densities were analyzed to quantitatively assess the reliability of the segmentation (Figure 5). The segmented mitochondrial volume data was also sub-divided into 100μm^3^ sub-volumes, and the mitochondrial densities were measured and tabulated (Figure 7).

**Figure 5.**
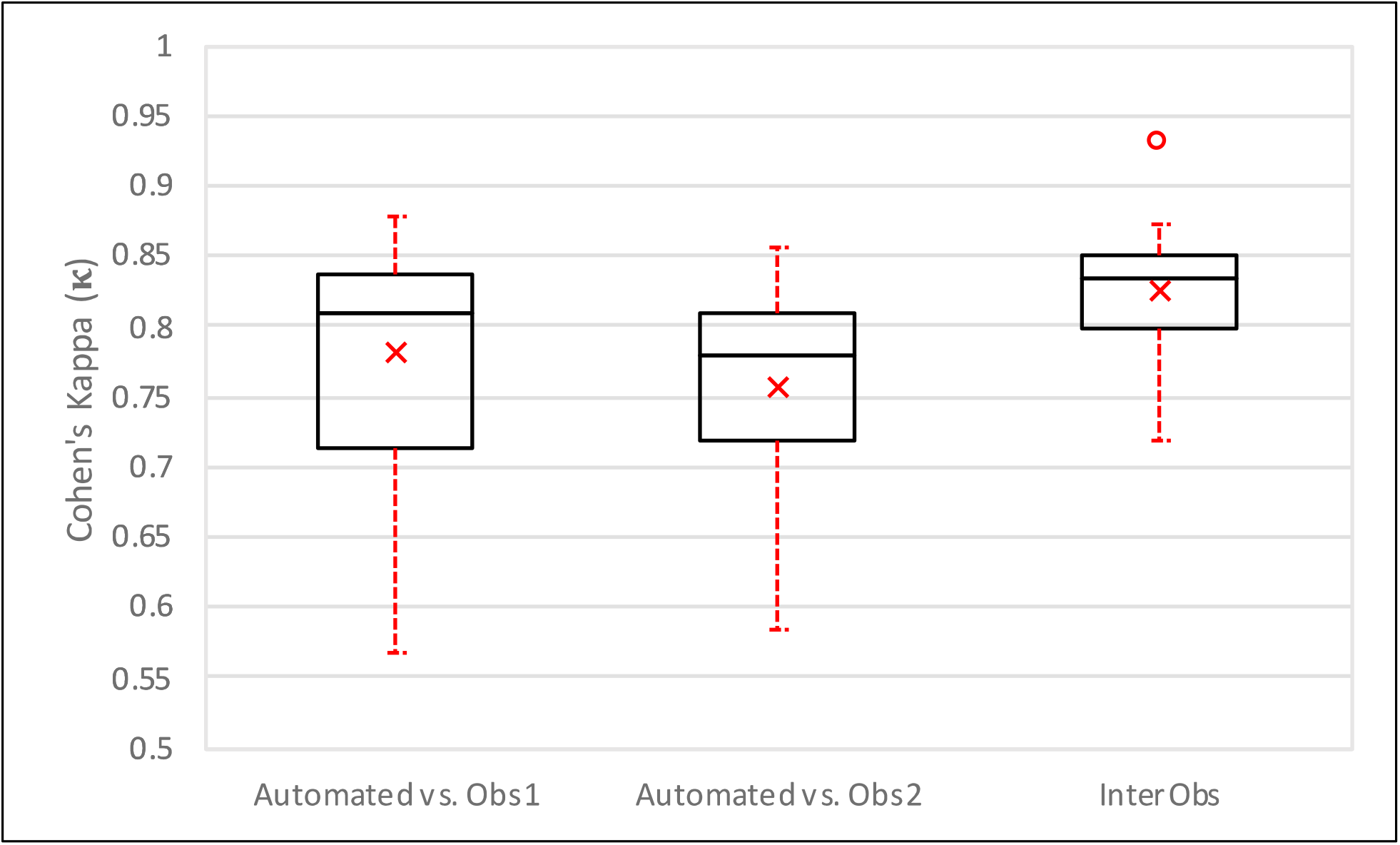
Boxplot of Quantitative Evaluation. Study of inter-observer variability and method versus each observer independently (n=12). The center line indicates the median value; a × indicates the mean, the box edges depict the 25^th^ and 75^th^ percentiles. The error bars show the extremes at 1.5 inter-quartile range, calculated inclusive of the median, excluding outliers, indicated by ∘.

Normalization of the average densities across different data sets minimized variability between data sets and allowed us to develop a generalized model of mitochondrial distribution across the muscle samples from different individuals. Combining multiple segmented volumes along each of the principal axes further increased the reproducibility of the results of automated segmentation.

### Quantitative Evaluation of Segmentation Pipeline against Manual Standards

Automated segmentation methods were compared to manually segmented volumes using the following metrics:

1. <u>Absolute Volume Difference (AVD) (%):</u> Absolute volume difference measurements were performed to measure the total volume difference between manually and automatically segmented volumes, allowing for a global metric of volume-to-volume difference. For labels with identical volume, %Difference (A, M) = 0, with increasing values indicating a greater volume difference between the two labels.
2. <u>3D Model-to-Model distance (Mean Surface Distance / MSD):</u> Both the manual and automated volumes were converted to ASCII mesh surfaces using ImageJ’s “3D viewer” (Chmid B 2010). These meshes were then transferred to the Cloud Compare platform (Cloud Compare 2018), where the manually segmented (reference) volume was compared to the automatically generated (comparison) volume, using the “Compute cloud/mesh distance” tool a map of the model-to-model distance was created. The max distance between the reference and automated datasets was set to 0.3 μm (any greater distance was set to the maximum threshold), and the model-to-model distance distribution was fitted to a Gaussian distribution, and the mean ± standard deviation calculated. A tricolor histogram was applied to the map with red representing automated density areas greater than manually segmented density, blue representing automated density areas less than manually segmented density and white represents <~15nm difference between structures.
3. <u>Sensitivity, Specificity, Accuracy, Dice Similarity Coefficient (DSC) and Cohen’s Kappa (κ) calculations.</u> Sensitivity, specificity and accuracy were calculated according to the conventional equations. The Dice similarity coefficient was calculated using a variation of the original formula (Dice 1945):

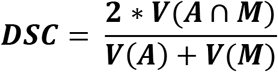

Cohen’s Kappa was calculated according to the equation found in (McHugh 2012):

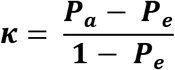

Where P_a_ = Actual Observed Agreement = **Accuracy**;

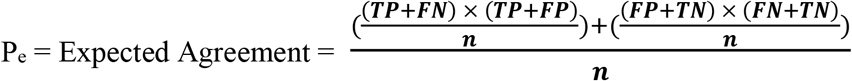

where n = total number of observations = **TP+FP+FN+TN**.

Where TP = True positive (**Automated [A] n Manual [M]**) ; FP = False Positive (**A\M**);

FN = False Negative (**M\A**) and TN = True Negative (**U\[AUB]**).

Table 1 shows the quantitative evaluation of the performance of the method vs two independently segmented versions of the same data set by two individuals along with inter-individual variability, calculated according to the equations above.

**Table 1:** Quantitative Evaluation. Study of inter-observer variability and method versus each observer independently (n=12), reported as a mean ± standard deviation.

|  | Automated vs. Observer 1 | Automated vs. Observer 2 | Inter-Observer |
| --- | --- | --- | --- |
| **Sensitivity (SENS)** | 0.9±0.07 | 0.92±0.06 | 0.83±0.12 |
| **Specificity (SPEC)** | 0.99±0.02 | 0.98±0.01 | 0.99±0.01 |
| **Accuracy (ACC)** | 0.98±0.02 | 0.98±0.02 | 0.99±0.01 |
| **Dice Similarity Coefficient (DSC)** | 0.79±0.07 | 0.79±0.08 | 0.83±0.05 |
| **Absolute Volume Difference (AVD) (%)** | 26.7±23.2 | 32.7±27 | 21.1±16 |
| **Cohen's Kappa ($\kappa$)** | 0.78±0.08 | 0.76±0.1 | 0.83±0.05 |

Sensitivity, specificity and accuracy are 0.83 ± 0.12, 0.99 ± 0.01 and 0.99 ± 0.01 respectively between the independent manually segmented data sets. A similar relative distribution of the mean sensitivity, specificity and accuracy were found between each manually segmented data set and the automated segmentation with values of 0.91 ± 0.07, 0.98 ± 0.01 and 0.98 ± 0.02 respectively. The relatively low sensitivity in all comparisons is indicative of the difficulty in defining the mitochondrial boundary, a 1-pixel difference in mitochondrial thickness across a volume can lead to dramatic decreases in sensitivity. However, there was excellent agreement between the two manual datasets and between the manual and automated datasets in the overall accuracy of identification of mitochondria, as illustrated using DSC and Cohen’s Kappa measurements, indicating a high level of agreement between the manually segmented data sets (0.83 ± 0.05 and 0.83 ± 0.05 respectively) and between each manual and automated segmented data (with average values of 0.79 ± 0.08 and 0.77 ± 0.1 respectively).

Figure 5 provides a graphical representation of Cohen’s Kappa values showing how the majority of the manual segmentations (75%) are above the widely accepted threshold of 0.7 for automated segmentations (McHugh 2012). We anticipate that this could be further improved with refinement of the classifiers or increasing the number of classifiers per volume.

Figure 6 provides a graphical representation of the model-to-model distance map between automated and manual segmentations of two muscle types. The mean surface distance (MSD) was calculated by fitting the above distributions (Figure 6C, 6D) to a Gaussian distribution, and the mean ± standard deviation was determined. The MSD showed the automated segmentation was accurate to 0.03 ± 0.06 μm, indicating a segmentation accurate to 2 or 3 voxels, with a slight bias to overestimate the size of the mitochondria relative to the manual segmentation. Of note, differences of the same magnitude were detected between observers, as mentioned previously and is indicative of the difficulty in defining precisely mitochondrial boundaries.

**Figure 6.**
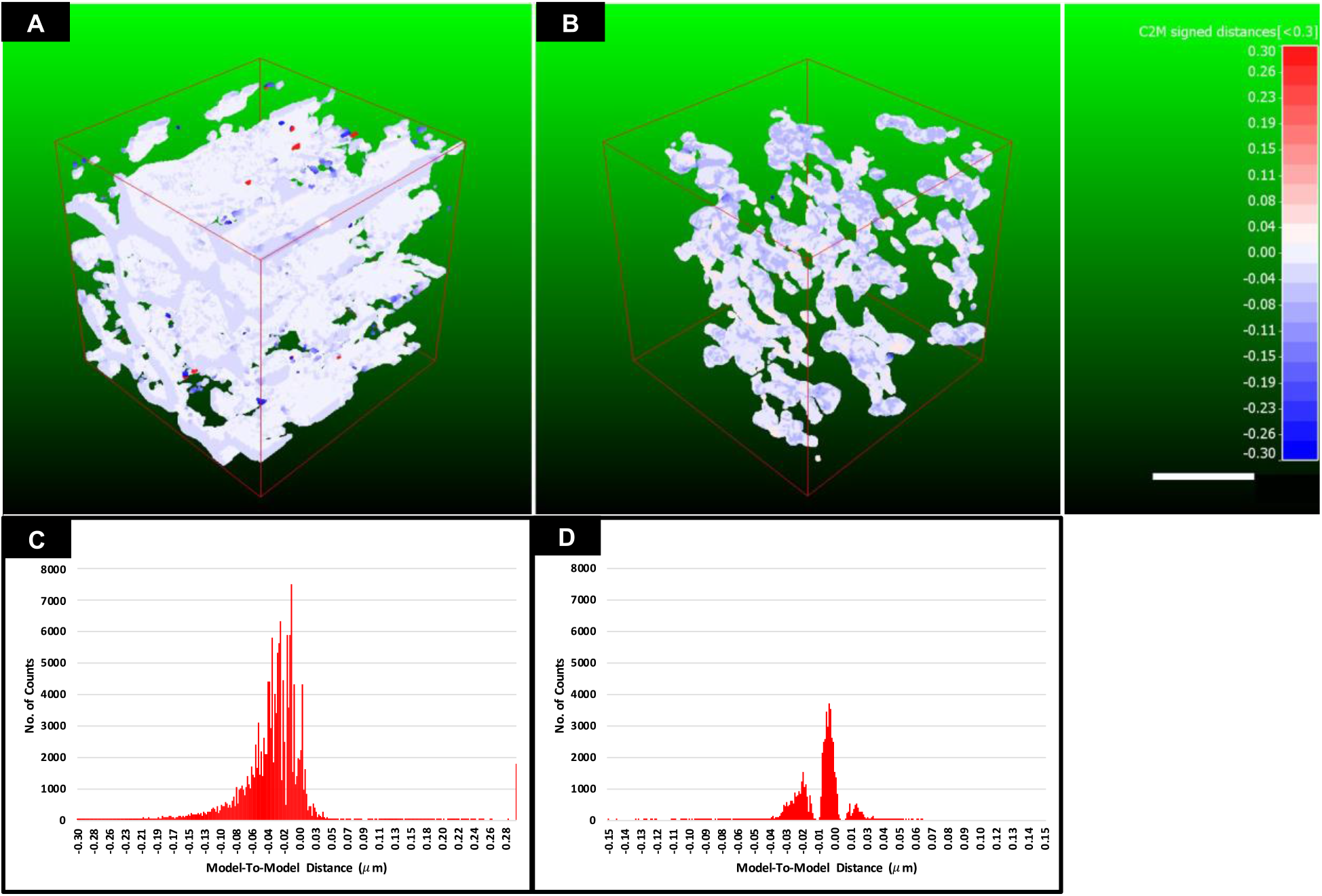
Model-to-Model Distance measurement. **A)** Isometric projection of 100 μm^3^ 3D model-to-model distance map for a typical “Type A” subvolume. **B)** Isometric projection of 100 μm^3^ 3D model-to-model distance map for a typical “Type B” sub-volume. **C)** A graphical representation of the distribution of the mean surface distances between manual (reference) and automated (comparison) volumes across the 3D mesh map for a typical type A sub-volume. **D)** A graphical representation of the distribution of the mean surface distances between manual (reference) and automated (comparison) volumes across the 3D mesh map for a typical type B sub-volume. Red-White-Blue distance map represents distances in microns, Red: Manual model > Automated model; White: Manual ≈ Automated model (± 15nm); Blue: Manual < Automated model. Scale bar = 2μm.

### Statistical Analysis of Mitochondrial Distribution in Human Skeletal Muscle

An essential step in the evaluation of this method was in determining its sensitivity to subtle differences in 3D volumes. Figure 7 demonstrates this by differentiating between two muscle types across 4 healthy individuals. This type of analysis has the potential to generate statistically relevant data for the study of age and disease-related differences in sub-cellular architecture across a population of individuals, where detection of subtle differences between populations may provide a wealth of insight into the mechanism and progression of disease states.

**Figure 7.**
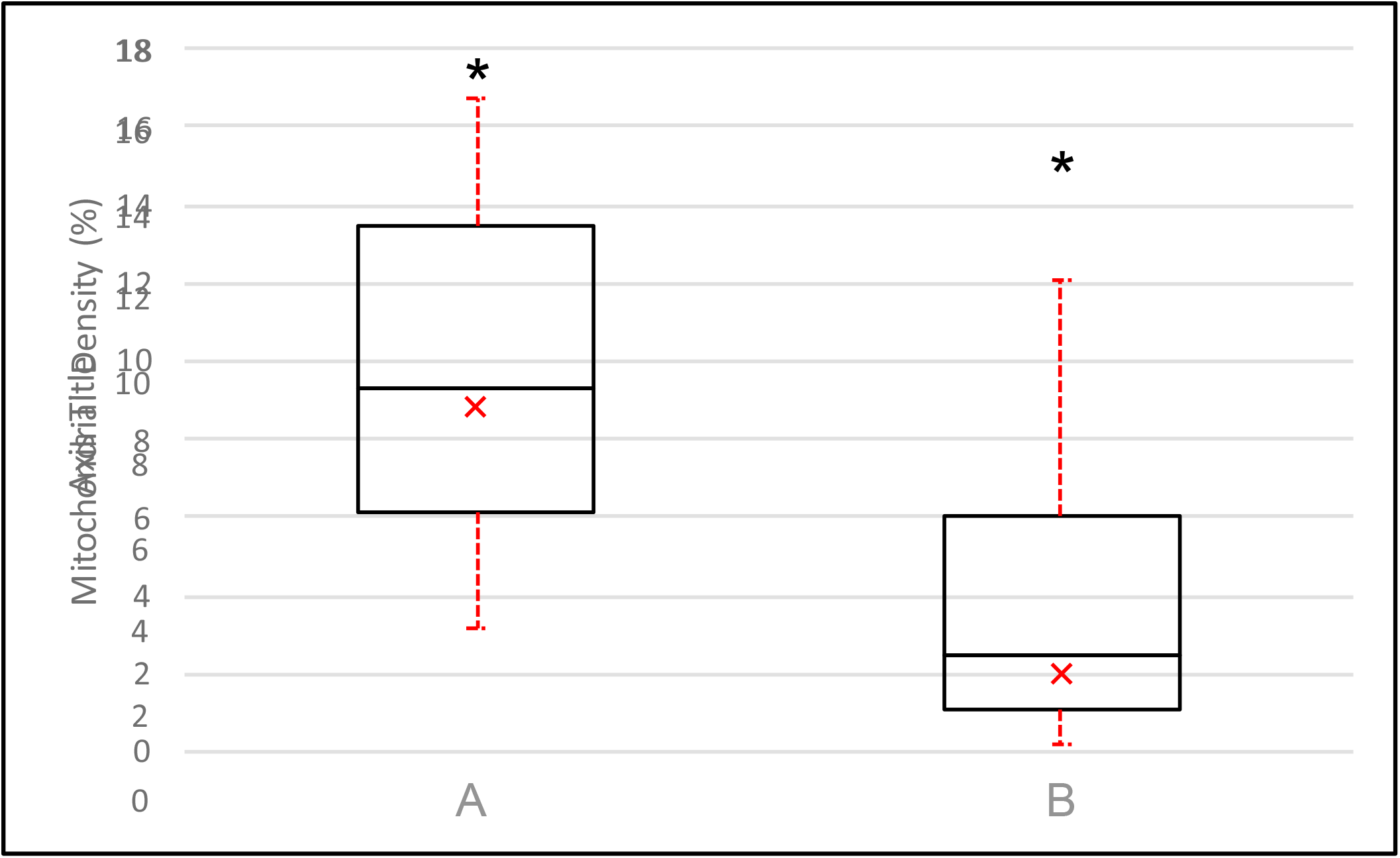
Boxplot graphical overview of mitochondrial distribution from two subpopulations of data (Type A vs Type B). The center line indicates the median values; a × indicates the mean, the box edges depict the 5^th^ and 95^th^ percentiles. The error bars show the maxima and minima of each population. * Indicates a statistically significant difference (p-value <<0.01,α = 0.05; Power (1β) = > 0.95). Total Sampled volume = 343,600μm^3^ across 4 healthy individuals.

### Evaluation of Automated Segmentation Performance using CA1 Hippocampal test dataset

Figure 8 is a demonstration of the performance of the segmentation approach against a hippocampal dataset. The time for obtaining this segmentation of 400μm^3^ volume took < 24 hours. We estimate that a 100-fold increase in the volume of the data to be segmented would not increase the segmentation time considerably, once the classes are produced they can be applied across an extremely large volume with little-added input.

**Figure 8.**
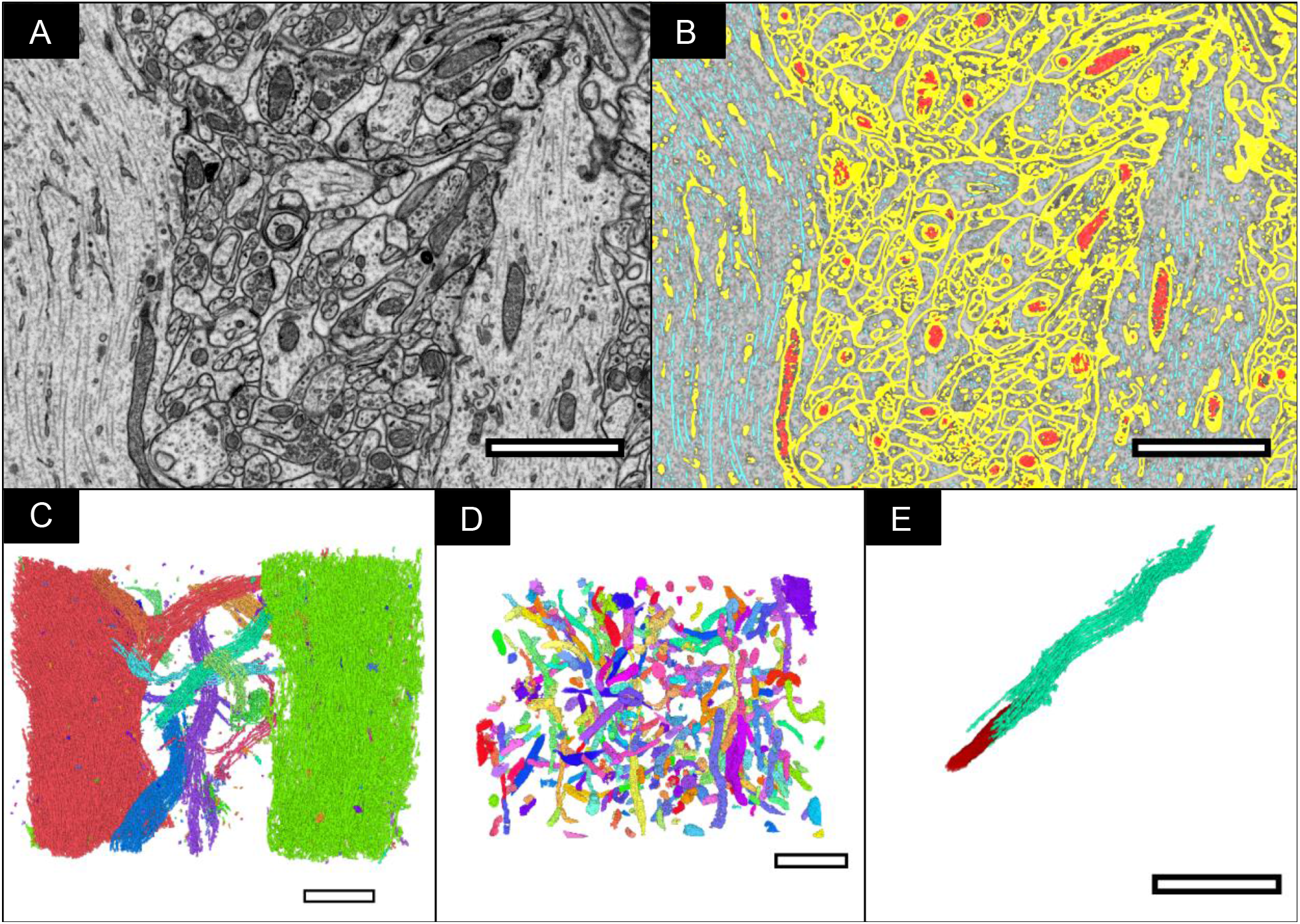
Other applications of this software. **A)** FIB-SEM slice from the CA1 hippocampal region of the brain with a voxel size of 5×5×5 nm^3^ **B)** FIB-SEM slice with automated segmentation overlaid (Yellow = Cell membrane ; Red = Mitochondria ; Cyan = Microtubules) **C)** 3D volume of segmented microtubules labelled separately, allowing for the straightforward isolation of individual cells for focused study. **D)** 3D volume of segmented mitochondria labelled separately. **E)** Individual microtubule (cyan) and mitochondria (red).

## DISCUSSION

In this work, we have presented a method for 3D segmentation and statistical analysis of human skeletal muscle volumes using an automated segmentation framework. The results demonstrate that rapid analysis of mitochondrial distribution in muscle architecture in relatively large volumes (>10,000 μm^3^) can be achieved consistently with high accuracy across multiple data sets. Our data collection approach enables rapid acquisition of large volumes at a rate of >1,500 μm^3^/hr. The acquisition rate is dependent on, among other variables, the pixel size which in turn determines the scanning area and resolution of subsequent volumes. Therefore, there is an inherent trade-off between resolution and volume acquisition rate. In this study, we determined that a 15nm^2^ pixel area returned a sufficient resolution and volume acquisition rate for statistical analysis of the mitochondria in the 3D image data from muscle tissue.

The Weka machine learning (Arganda-Carreras, et al. 2017) software was chosen specifically for its segmentation capabilities. The Weka software is a robust software that is professionally maintained by The University of Waikato in New Zealand. Weka’s software is powerful and versatile, allowing it to be ported to various operating systems and be used as a component of larger software. Our approach to full volume segmentation is to manually classify a small set of images and then export the manually trained classifier to use on the entire data set. These methods are generalizable to a variety of other data sets.

Large-scale, high-resolution volume segmentation and validation of multiple cellular components can be achieved by a single individual in an extremely brief timespan using our approach. We illustrate this using a publicly available dataset (Computer Vision Laboratory - Electron Microscopy Dataset 2018) used as a standard to test automated segmentation approaches. This dataset was acquired and segmented at a spatial resolution of 5nm^3^ and produced several 3D segmentations of major cellular organelles in less than 24 hours. Currently, the majority of neuronal tissue segmentations (Zheng, et al. 2018) are performed using manual tracing methods, however, due to its time-consuming nature, much of the intra-cellular detail is lost. Through the use of our approach, this information can be rescued and used in conjunction with manually traced data to build a complete picture of the sub-cellular environment in neuronal tissues. In conclusion, we note that our approach, which is available online to any interested user, can be readily applied to a wide variety of biological problems, with minimal human input, from tackling large-scale population-wide studies to the sensitive high-resolution analysis of cellular components.

## Supporting information

Supplementary Guide

## MATERIALS AND METHODS

### Candidate Selection and Muscle Biopsy

This study was conducted in healthy men participating in the Baltimore Longitudinal Study and Aging (BLSA) and the Genetic and Epigenetic Signatures of Translational Aging Laboratory Testing (GESTALT) studies. The design and description of the BLSA and GESTALT studies have been previously reported (Tanaka, et al. 2018; Shock, Greulich and Andres 1984; Stone and Norris 1966). Skeletal muscle biopsies were performed in fasting conditions as described elsewhere (Gonzalez-Freire, et al. 2018). Briefly, a ~ 250mg muscle biopsy was obtained from the middle portion of the vastus lateralis muscle using a 6-mm Bergstrom biopsy needle inserted through the skin in the muscle. A small portion of muscle tissue (~5mg) was immediately placed in 2% Glutaraldehyde (GA) and 2% Paraformaldehyde (PFA) in 100mM sodium cacodylate buffer, pH 7.3-7.4 at 4°C until required for sample preparation. The rest of the biopsy specimen was snap frozen in liquid nitrogen and subsequently stored at −80°C until used for further analyses.

### Fixation, Contrasting, and Embedding

Muscle biopsy samples from human donors were fixed with 5% glutaraldehyde in 100mM sodium cacodylate buffer at pH 7.4 as in a murine skeletal muscle study (Glancy, Hartnell and Malide, et al. 2015). In order to achieve the contrast required to be able to consistently identify mitochondria with similar signal to noise ratio the standard post-fixation protocol used for the murine muscle skeletal muscle samples was changed. Here we post-fixed with 2% Osmium Tetroxide (OsO4) in sodium cacodylate buffer for 1hr at RT, washed with ddH2O and treated with 4% tannic acid in sodium cacodylate buffer. A second treatment of 2% OsO4 in cacodylate buffer either reduced or not reduced with 0.6% Potassium Ferrocyanide was performed for 1hr at RT. Samples were then washed in ddH2O and treated with 2% Uranyl Acetate (UA) in ddH2O at 4°C overnight (Kobayashi, Gunji and Wakita 1980; Lewinson 1989). The samples were then washed in ddH2O, 5 x 10 min, and dehydrated using a graded ethanol series ending in 100% propylene oxide. Infiltration of embedding media was performed using a ratio of 2:1, 1:1, 1:2 propylene oxide to Eponate12 resin formula (EMS). Samples were embedded in resin molds and placed in an oven set at 60°C overnight for polymerization.

### Area selection for FIB-SEM analysis

Areas of muscle were chosen for FIB-SEM data collection following a survey of 0.5-1 μm thick sections of resin-embedded muscle tissue; sections were created using an Ultracut S microtome from Leica Microsystems. The sections were stained with Toluidine blue which stains nucleic acids blue and polysaccharides purple. Once stained, the orientation and morphology of the fiber was assessed using a light microscope. Suitable areas with intact muscle fibers were chosen for FIB-SEM data collection using the last section taken from the top of the block-face, and digital images were taken for reference. These images were used as maps to pinpoint the previously selected areas for data collection in the FIB-SEM (Glancy, Hartnell and Malide, et al. 2015). The resin was then cut to create a suitable sample for SEM. The samples were then sonicated in ethanol: water (70:30) for 15 mins to remove dust and particulates which would hinder imaging. The sample was then mounted on an aluminum stub using a double-sided adhesive conductive carbon tab, and the sides painted with silver paint to prevent charge build-up.

The sample was then allowed to dry, placed in a sputter coater (Cressington model 108), and coated with gold for 40 seconds at 30 mA.

After gold coating, the sample was placed into the sample chamber of the FIB-SEM. FIB-SEM imaging was performed using a Zeiss NVision 40 microscope, with the SEM operated at 1.5 keV landing energy, a 60 μm aperture and backscattered electrons were recorded at an energy selective back-scattered electron (EsB) detector. The user interface employed ATLAS 3D from Carl Zeiss, consisting of a dual 16-bit scan generator assembly to simultaneously control both the FIB and SEM beams and dual signal acquisition inputs, as well as the necessary software and firmware to control the system.

The fiber of interest was located using the SEM, and the instrument was then brought to eucentric and coincidence point at a specimen tilt of 54°, i.e. the specimen height where the specimen does not move laterally with a change in tilt and where the focal point of both FIB and SEM coincide. Once the exact milling area was determined with reference to the microscope images, a protective platinum pad was laid down on top of the area using a Gas Injection System (GIS) of size 60 μm x 30 μm and 5 μm in thickness. Then alignment marks were etched into the platinum pad using an 80 pA FIB aperture to allow for automated tracking of milling progress, SEM focus and stigmation during acquisition. After alignment etching, the platinum pad was covered with a carbon pad using the GIS to protect the etched marks from the milling process. After deposition of the carbon pad, a trench was dug using a 27nA FIB aperture to allow for line-of-sight for the SEM ESB detector. After the trench was dug, the imaging face was polished using a 13 nA FIB aperture. The FIB aperture was changed to 700 pA and SEM imaging area selected (Typical Image size: 4000 px x 2000 px /Pixel size: 15 x 15 x 15 nm [xyz]) the automated acquisition software was set up and run until all the sample area was acquired.

### Image processing and segmentation

After SEM acquisition the individual image files (.tif) were aligned using a cross-correlation algorithm (Murphy, et al. 2011). The images were then opened in ImageJ, and the volume was cropped to ensure a minimum distance of at least 1 μm away from the cell boundary in any direction, this was performed to reduce measurement variability of mitochondrial density due to the non-uniform distribution of mitochondria near capillaries and cell boundaries. (Sjöström, et al. 1982)

To reduce noise volumes were median filtered by 1 pixel in the x, y and z directions and then binned by 2 in all three axes to produce a final voxel size of 30 x 30 x 30 nm. The volume’s contrast was normalized and equalized using ImageJ’s “Enhance Contrast” function.

Sample images were required for preliminary training to construct the necessary machine learning classifiers for automatic segmentation. Referring to the schematic in Figure 10, three representative slices (one from each 3 ^rd^ of the volume), were taken at random from each of the principal axes: x, y and z (9 slices in total) (Figure 10A). A classifier was trained for each slice by sampling several main structures found in each sample image (Figure 10B). The primary structures, based on standard histological examples, were as follows: z-disk, mitochondria, A-band, I-band, sarcoplasmic reticulum and lipids. The classifier was trained using all training features available in the “Trainable Weka Segmentation” plugin for ImageJ Fiji, a robust machine learning plugin that is professionally maintained by The University of Waikato. The Weka algorithm, in brief, extracts image features using common filters that can be categorized as edge detectors (e.g. Laplacian and Sobel filters), texture filters, (such as minimum, maximum, and median filters), noise reduction filters (such as Gaussian blur and bilateral filter), and membrane detectors, which detect membrane-like structures of a specified thickness and size. Furthermore, the feature set also included additional features from the ImageScience suite (https://imagescience.org/meiiering/software/imagescience).

Since only 2D image features were calculated, classifiers were trained and applied on all three image axes to compensate for the loss of a third dimension. In our machine learning approach, we applied the multi-threaded version of the random forest classifier with 200 trees and 2 random features per node. Probability measurements of each class were generated, allowing for a class-by-class assessment of the performance of each classifier during training. Segmentation masks of the key skeletal structures were then outputted based on these probability measurements (Figure 9).

**Figure 9.**
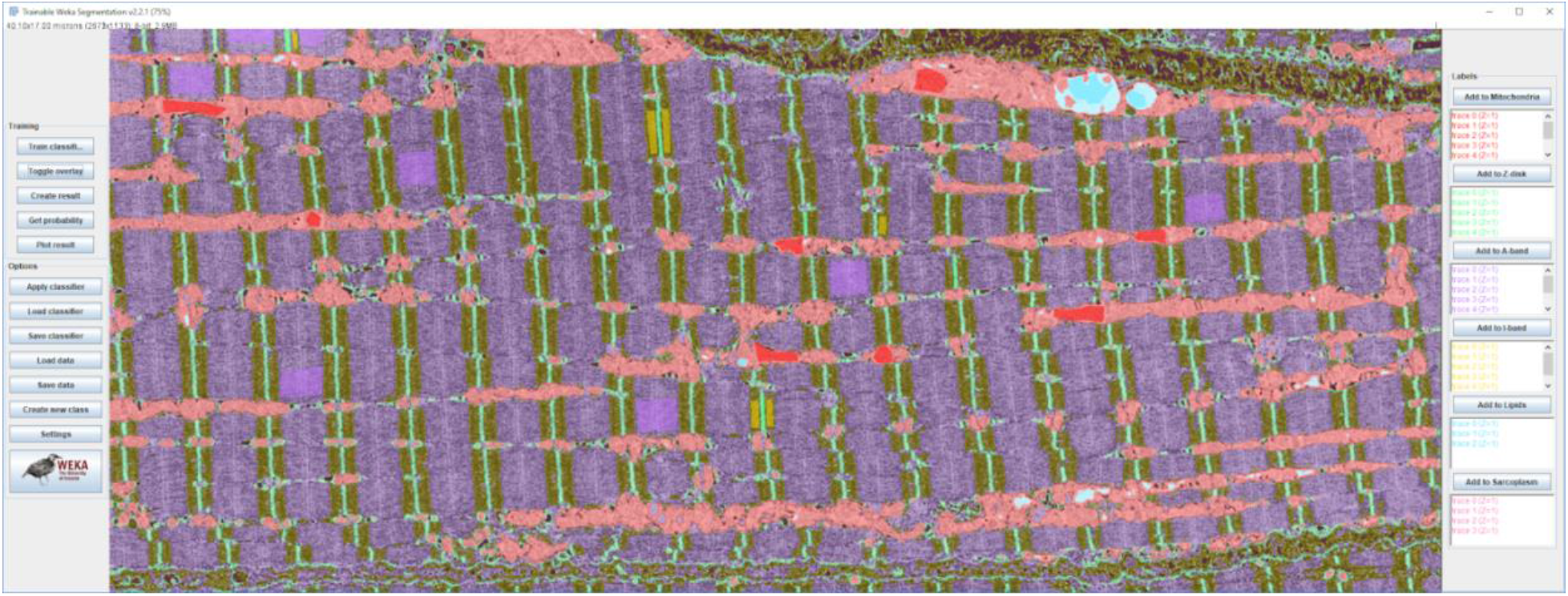
Example of Weka software manual classification in ImageJ.

Once all 9 classifiers were trained (3 for each axis), they were exported as separate “.model” files and applied to each slice in the volume according to the respective axes which they were trained.

The segmentation of the full volume was performed on the Biowulf supercomputer cluster, and its implementation is as follows:

To simultaneously process many image slices at once, each slice was opened in a separate instance of ImageJ Fiji, and then Weka machine learning was executed in each instance. Since only 2D image features were calculated, each instance of ImageJ Fiji could effectively classify an image without needing access to any other image data.

Thus, images were simultaneously classified by parallel processors running multiple instances of ImageJ Fiji. Each instance executed a Beanshell script (source.bsh) that automatically performed Weka machine learning on a specified image using a specific classifier file. The process of opening instances of ImageJ Fiji was automated through the command line interface by using the existing “–headless” option that came with the ImageJ Fiji package. Biowulf effectively allocated and launched hundreds of processors at once with the use of the “swarm” command that already existed on the supercluster.

The command required a formatted file containing independent commands to distribute to each processor and to generate such a file quickly we wrote an automated Bash script “generate_swarm_script.sh”. If a different system other than Biowulf is being used, then it is advised to create a script that launches parallel instances of ImageJ Fiji that execute the “source.bsh” Beanshell script. It should be noted that Weka machine learning is optimized to run faster by utilizing a substantial amount of RAM. For the classification of our large FIB-SEM images, we allocated 25 GB RAM per processor per image.

A total of 9 automated segmentation volumes were created. The 3 volumes from each axis were first added together using ImageJ’s “Image calculator” function, and density which did not overlap with at least one of the other 2 volumes was removed through simple thresholding (Figure 10C). Each axis volume was then added together using the previously mentioned function, and density which did not overlap with at least one of the other 2 axes was removed through simple thresholding (Figure 10D). The volume was then filtered by 2 pixels in the x, y and z directions using ImageJ’s “Median 3D Filter” function and was used for statistical analysis of mitochondrial densities.

**Figure 10.**
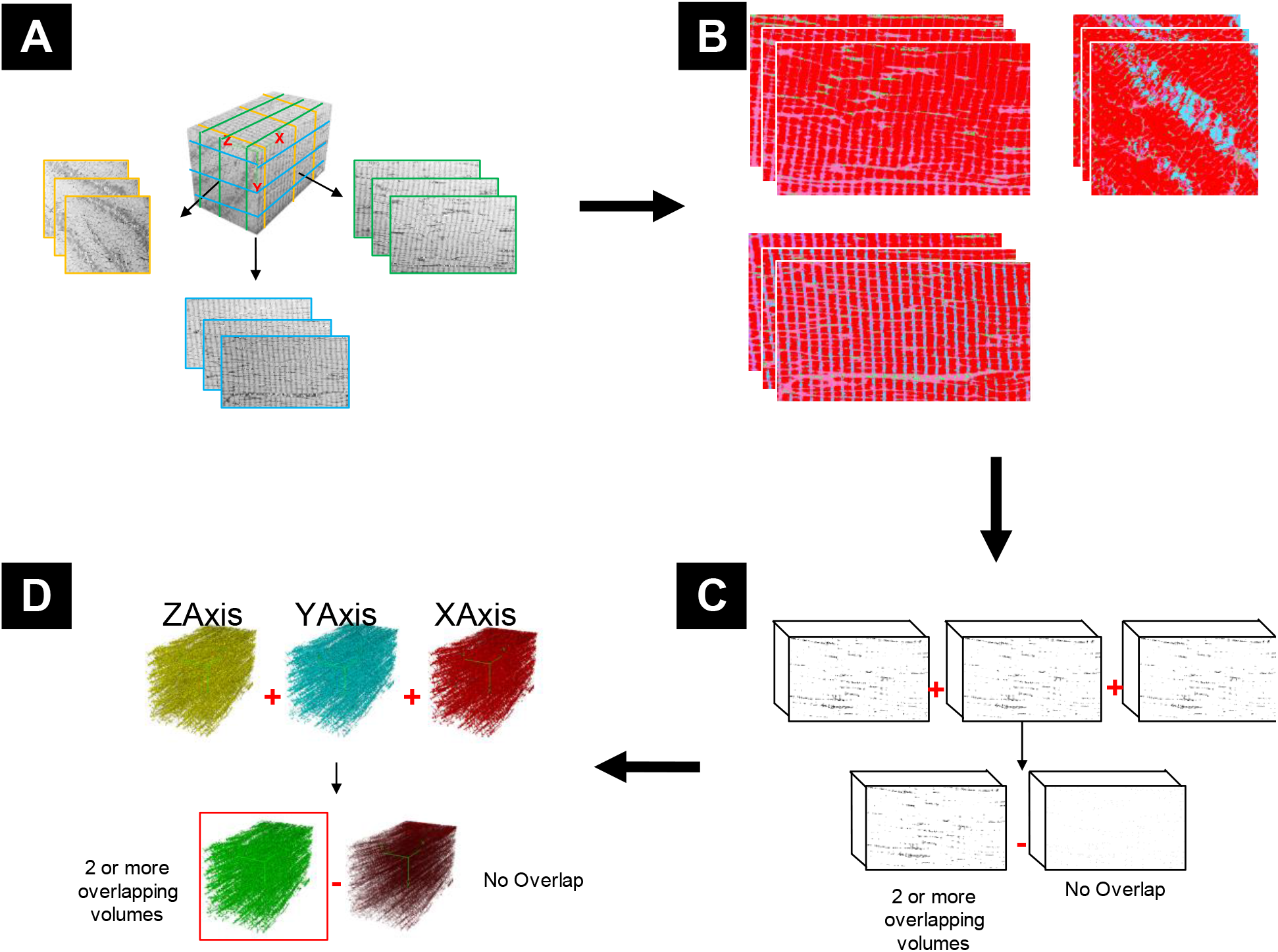
Graphical representation of key steps in segmentation pipeline. **A) Classifier Selection**: Three representative slices are taken from each of the principal axes. **B) Classifier Generation**: Each slice is manually classified based on the organelles within the volume. **C) Volume Classification and Refinement**: The classifiers are applied to the entire volume and produce segmented volumes of each class. The mitochondrial class is isolated, and each of the 3 volumes from the same axes are combined and non-overlapping data removed to produce an axial volume. **D) Axial Volume Combination**: Each of the refined volumes from the principal axes are combined, and non-overlapping data is removed.

## ACKNOWLEDGMENTS

We thank members of the Ferrucci and Subramaniam laboratories for helpful discussions. We thank Kunio Nagashima for resin infiltration and embedding of muscle samples. We thank Jessica De Andrade for help with validation of the automated segmentation. This research was supported by the Intramural Research Programs of the National Institute of Aging, NIH, Baltimore, MD to LF, by funds from the Center for Cancer Research, National Cancer Institute, NIH, Bethesda, MD to SS. This work utilized the computational resources of the NIH HPC Biowulf cluster (http://hpc.nih.gov).

## Author Contributions

BC: FIB-SEM data collection, data processing, ML training, data interpretation, method validation, paper writing

AM: Generated ML platform cloud computing capabilities, paper and user guide writing

MGF: Candidate selection, sample biopsy

LH: FIB-SEM data collection, data interpretation

LF: project design, data interpretation

SS: project design, data interpretation, paper writing

## SUPPLEMENTARY MATERIAL

**Figure 2. - Figure Supplement 1.**
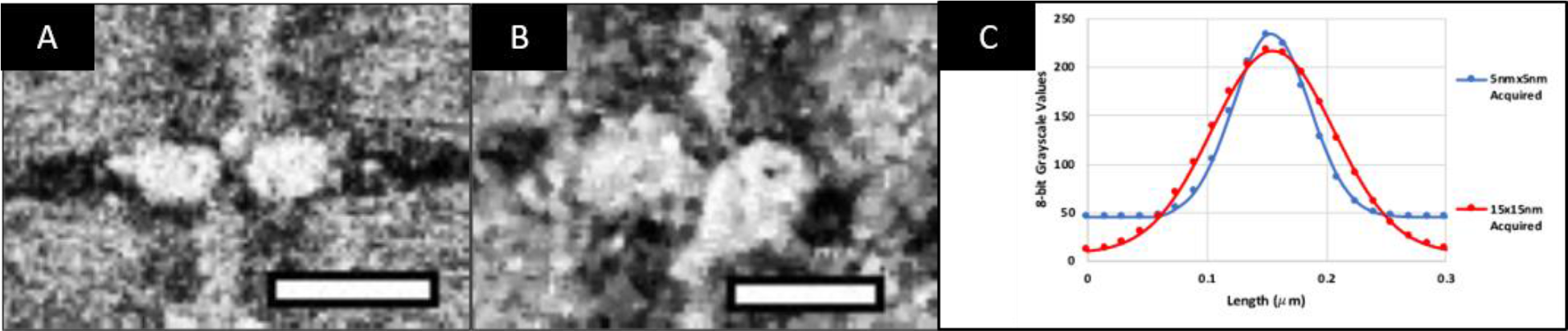
Resolution Comparison. **A)** Slice of normal skeletal mitochondria acquired at 5×5nm (xy) pixel size and binned by 3 to 15×15nm pixel size. **B)** Slice of normal skeletal mitochondria acquired at 15×15nm pixel size. **C)** A comparison of the difference between mitochondria in image A and B, indicating an increase in the signal-to-noise ratio (SNR) and a ~10% loss in resolution i.e. Full Width at Half Maximum (FWHM) between the sample acquired at 15×15nm relative to the sample acquired at 5×5nm. Scale bar = 500nm

**Figure 3 - Figure Supplement 1. Video of Automated Segmentation of Type A muscle**

See attached supplementary file

**Figure 3 - Figure Supplement 2. Video of Automated Segmentation of Type B muscle**

See attached supplementary file

**Figure 8. - Figure Supplement 1. Video of Automated Segmentation of CA1 Hippocampal tissue**

See attached supplementary file

## Abbreviations

FIB-SEM: Focused Ion Beam-Scanning Electron Microscopy
ML: Machine Learning
AVD: Absolute Volume Difference
MSD: Mean Surface Distance
DSC: Dice Similarity Coefficient.

