## Supplementary Guide for "Automated methods for 3D Segmentation of Focused Ion Beam-Scanning Electron Microscopic Images"

### GUIDE TO BULK SEGMENTATION OF FIB-SEM IMAGES USING HIGH PERFORMANCE MACHINE LEARNING

#### Introduction

This method was developed by Alexander V. Maltsev and Brian Caffrey as part of their work in a joint collaborative effort between the Translational Gerontology Branch at IRP/NIA/NIH and the Laboratory of Cell Biology at IRP/NCI/NIH. The program uses ImageJ Fiji, Weka machine learning, and various programming scripts to rapidly and automatically segment structures in FIB-SEM images of human skeletal muscle. This program is still in its beta stage and its functionality so far has only been tested on human skeletal muscle and the CA1 hippocampal region of the brain.

Since the program was created at the NIH, a US Government agency, it belongs to the public domain and is allowed to be freely distributed for the purpose of improving healthcare in the US and worldwide. In this research paper, we provide a detailed methodology on how to setup and use our software pipeline. If you use or further develop our software, please credit our work by citing our research paper.

#### Setting up Software for Preprocessing and for Classifier Training

1. To perform necessary image preprocessing, download the ImageJ Fiji software from: https://fiji.sc/. Make sure the distribution type matches the operating system of your computer.
2. Check to make sure the following jar files are the necessary versions:
   a) plugins/Trainable_Segmentation-3.2.11.jar
   b) jars/ij-1.51n.jar,
   c) jars/fiji-lib-2.1.1.jar,
   d) plugins/Anisotropic_Diffusion_2D-2.0.0.jar,
   e) jars/VIB-lib-2.1.1.jar,
   f) jars/commons-math3-3.6.1.jar,
   g) jars/weka-dev-3.9.0.jar,
   h) jars/imglib2-algorithm-0.6.2.jar,
   i) jars/imglib2-algorithm-gpl-0.1.5.jar,
   j) jars/imglib2-ij-2.0.0-beta-35.jar,
   k) jars/imglib2-3.2.1.jar,
   l) plugins/Stitching_-3.1.1.jar
3. To run Weka machine learning classification with additional features, download the jar files named FeatureJ_.jar and imagescience.jar from: https://imagescience.org/meijering/software/featurej/ and then transfer them to the "plugins" folder under the ImageJ Fiji directory named "Fiji.app".

#### Preprocessing Images

1. Launch the ImageJ Fiji software on a desktop computer.
2. Place into a folder data to be segmented and then drag the folder into ImageJ Fiji menu bar.
3. A pop-up will ask whether or not to open the images as stack. Click yes. The images should now be opened in ImageJ Fiji as an image series in a single spatial plane.

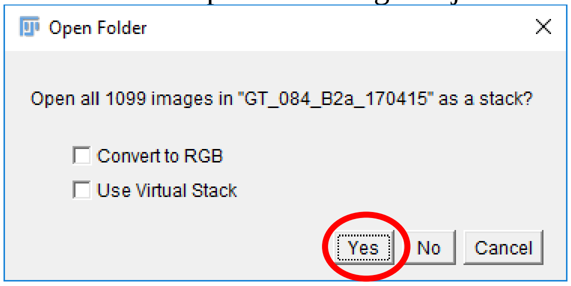

4. Click on the stack and normalize it using Process -> Enhance Contrast. Check "Normalize" and "Equalize Histogram" and click Ok.

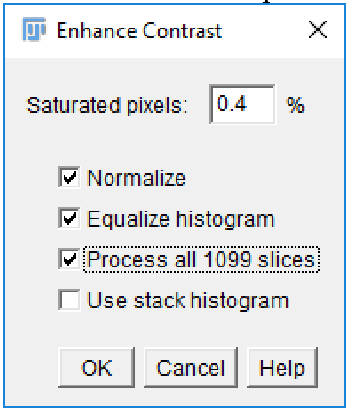


5. Sample 3 slices from the sequence, each one about a third into the sequence using the option Image -> Duplicate. Specify the image number in the textbox to the right of "Range".

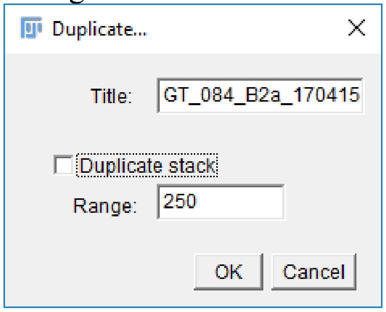


6. Export the normalized and equalized stack as an Image Sequence to a new folder using the option File -> Save As -> Image Sequence. Make sure the image names do not have any special characters, only letters, numbers, and dashes.
7. Project the opened image sequence onto the XZ spatial plane using the option Image -> Stacks -> Reslice and in the window that pops up, select the option "Top" next to the label "Start at".

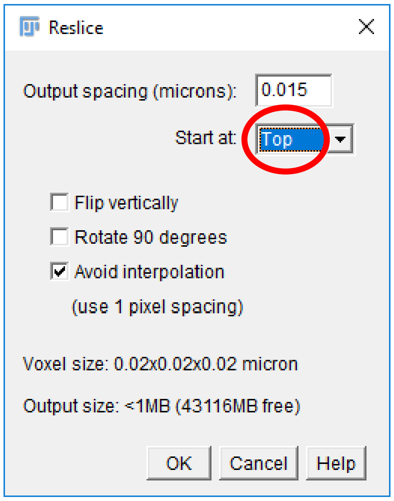

8. Repeat steps 4-6 for the XZ projection of the image sequence.
9. Project the opened image sequence onto the YZ spatial plane using the option Image -> Stacks -> Reslice and in the window that pops up, select the option “Right” (or “Left”) next to the label “Start at”.

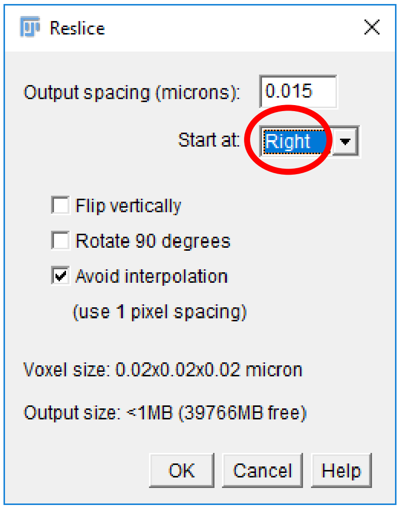


10. Repeat steps 4-6 for the YZ projection of the image sequence.
11. As a result of preprocessing, there should be 3 exported image sequences of each projection (XY, XZ, YZ), each sequence completely normalized and equalized. Furthermore there should also be 9 total sampled images, 3 from each projection.

**Training Classifiers for Machine Learning**

1. Input the image samples, one at a time, into the Weka Trainable Segmentation in ImageJ Fiji with the option Plugins -> Segmentation -> Trainable Weka Segmentation.
2. Make the necessary amount of classes for segmentation. We used the classes: mitochondria, z-disk, A-band, I-band, lipids, and sarcoplasm. Make new classes using “Create new class” button and modify the class names in the Settings.

3. To check whether all training features are available, click Settings and all checkboxes should be available to use. While you're at the Settings, check all boxes.

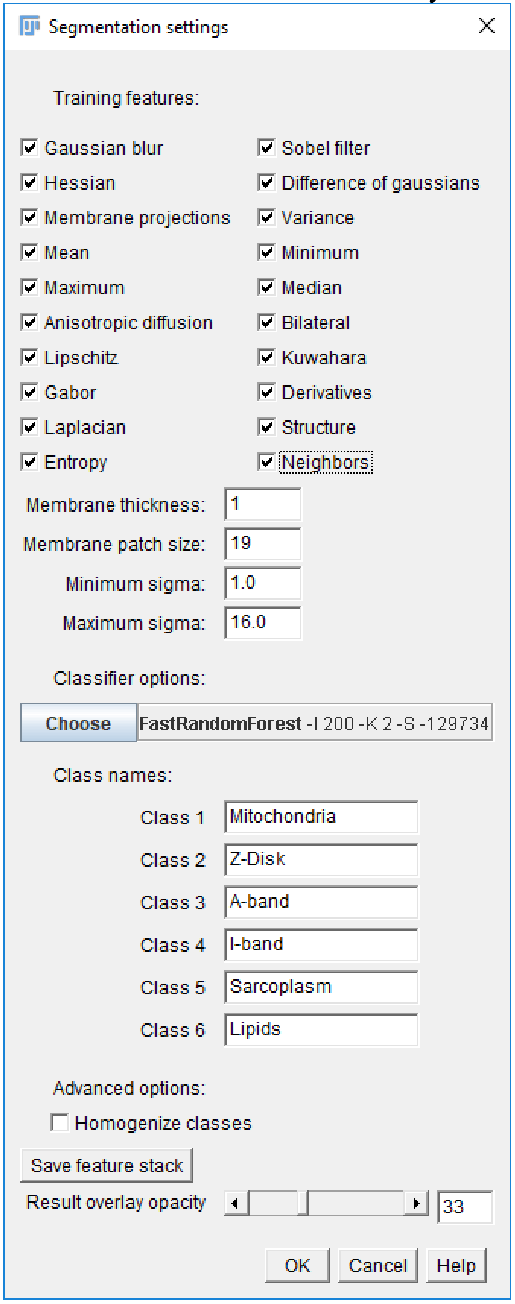

4. Click on the label FastRandomForest and set options as shown (note that the value 0 in some areas lets the classifier know to automatically set the largest value possible for the dataset and the hardware).

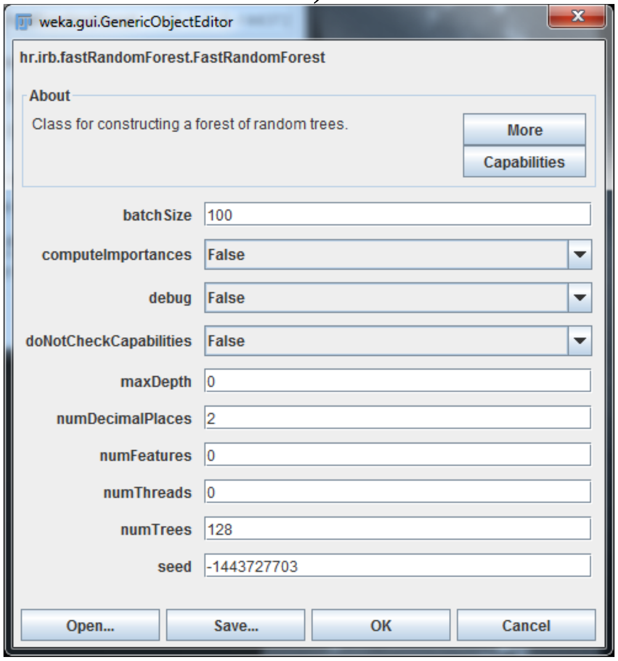


5. To train a classifier, first begin by sampling areas for each class using the selections in the ImageJ Fiji menu bar. Polygon selection is the most flexible and recommended way to sample. Make sure that the sample does not include any surrounding area. The accuracy of the outcome will be higher with more samples taken. The image can be zoomed into by using ctrl-(scrolling with a computer mouse) and moved around with using the hand icon in the menu options. Classifiers can be retrained endlessly to achieve desired classification.

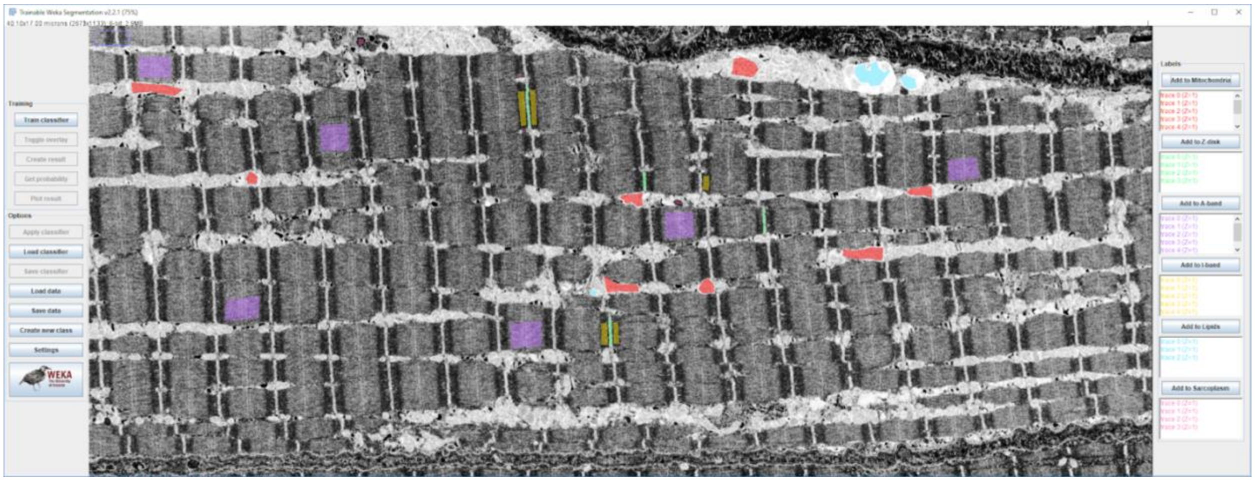

6. After sampling is complete, train the classifier using the “Train Classifier” button.

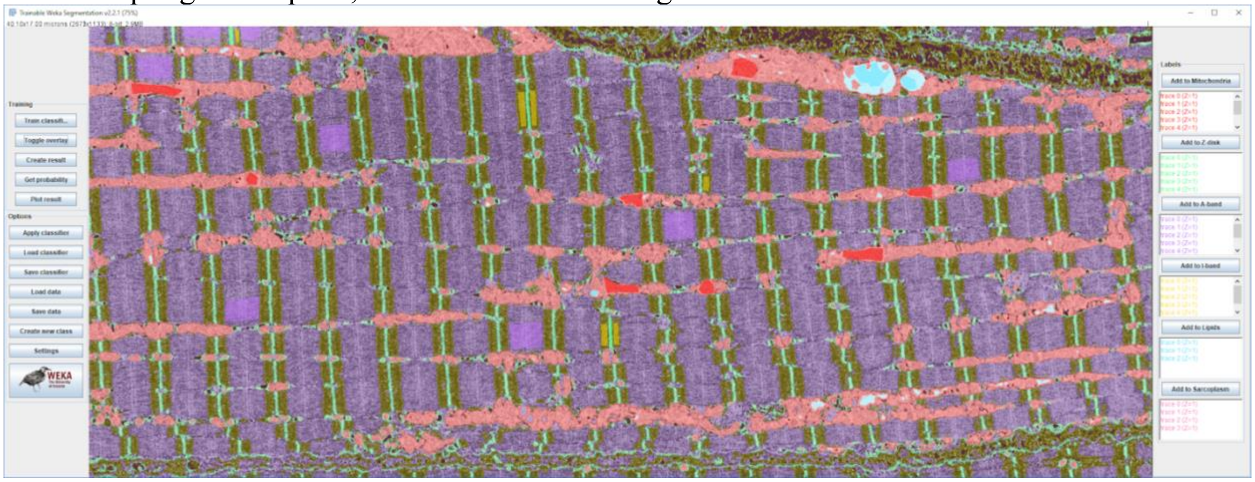


7. After the result is obtained, if necessary, sample again to retrain the classifier so that obvious misclassifications can be fixed. This retraining should be faster than the first training. Retrain the images until classification is satisfactory.
8. Save the classifier using the button “Save Classifier”. Make sure the saved classifier name does not have any special characters, only letters, numbers, and dashes.
9. Repeat steps 1-8 for all 9 image slice samples.
10. As a result of steps 1-9, there should be 9 exported classifier files (.model), 3 for each spatial projection (XY, XZ, YZ).

#### Performing High Performance Machine Learning on server

1. To perform high performance Weka machine learning on server, make sure all scripts and program files are on your system, including fiji.

2. Move the 9 generated classifiers from classifier training to the directory that holds the "ImageJ-linux64" executable file using MobaXterm. Also transfer the 3 normalized and equalized image sequences of each projection (XY, XZ, YZ) to your server. The image sequences do not have to be inside the "Fiji.app" directory. The final collection of files should look like the figure below.

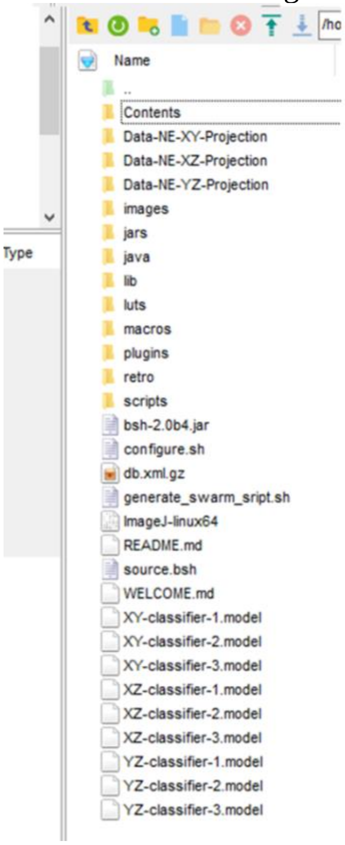


3. Create a new folder on the server that will hold the resulting segmentation maps. The folder does not have to be inside the "Fiji.app" directory.
4. Navigate to the directory that holds "generate_swarm_script.sh". Execute the command "./generate_swarm_script.sh [ INPUT FOLDER ] [ CLASSIFIER MODEL ] [ RESULT MODE ] [ OUTPUT FOLDER]", filling in necessary parameters and removing square parenthesis. Specify the full path of each file and folder and make sure to match each classifier with the correct projection that it was trained on. The RESULT MODE parameter can either be "Probabilities" or "Segmentation". For our purposes, we used the option "Segmentation" which automatically classified pixels based on which class they most likely belonged to. After executing the "generate_swarm_script.sh" script, a formatted swarm file named "run.swarm" should appear in the same directory. If on-the-fly explanation is needed, execute the command "./generate_swarm_script.sh --help" for immediate details on parameters and options.
5. To perform machine learning on images, execute the command "swarm –f run.swarm –t 1 –p 2 –g 25". The parameters allocate 25 GB of RAM and 1 thread for each processor to classify a single image slice. The swarm script used in this step can be found on github, here.

6. Once no more jobs with the swarm number are executing, then the classification is complete and the resulting probability maps should be in the OUTPUT FOLDER specified.
7. Repeat steps 4-7 with the next classifier (out of 9) and make sure to enter a new OUTPUT FOLDER to keep results differentiable and organized. The "swarm" command can be launched simultaneously with other "swarm" commands to maximize performance.
8. When step 8 is complete with all 9 classifiers, 9 sets of segmentations should be generated, each in a separate folder. Check to make sure all images have been processed. If some images fail to segment or are missing, place the missed images in a new folder and redo steps 5-8 on those images.
9. Once all images have succeeded processing, transfer using MobaXterm, all 9 folders that hold a segmentation result to a Desktop computer.

**Post Processing to Form the Final Segmentation Volume**

1. Launch the ImageJ Fiji software on a desktop computer.
2. Import three segmented volumes at a time into ImageJ Fiji that correspond to each spatial dimension. This is done by dragging and dropping each folder into the ImageJ Fiji menu bar.
3. Convert each stack from 32-bit to 8-bit with the option Image -> Type -> 8-bit
4. Invert each stack using Edit -> Invert
5. Threshold each result to include only the segmented mitochondria with the option Image -> Adjust -> Brightness/Contrast and place the threshold bar in the middle of the ticks. The figure below shows the settings to displays pixels with 3 overlaps present (this ensures maximum consensus between the different segmented volumes).

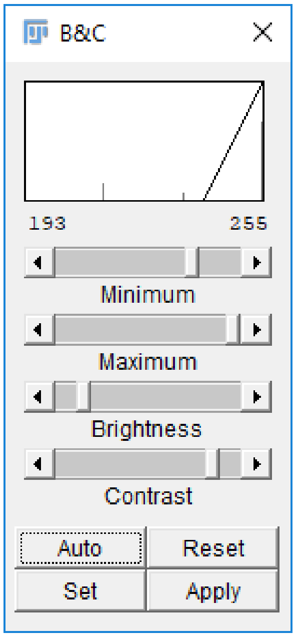

6. From the “Preprocessing” section, do “Reslice” to make all the 3 models in the same projection. Use the XY projection as the base, then reslice the XZ projection using the parameter “Top” and then reslice the YZ projection using the parameter “Top” and rotate 90 degrees.
7. Rename the resulting Resliced projections to differentiate between the segmentation results.
8. Add each volume mask together using the option Process -> Image Calculator. After adding 2 of them together, convert from 32-bit to 8-bit. After adding the 3rd mask, convert from 32-bit to 8-bit again.

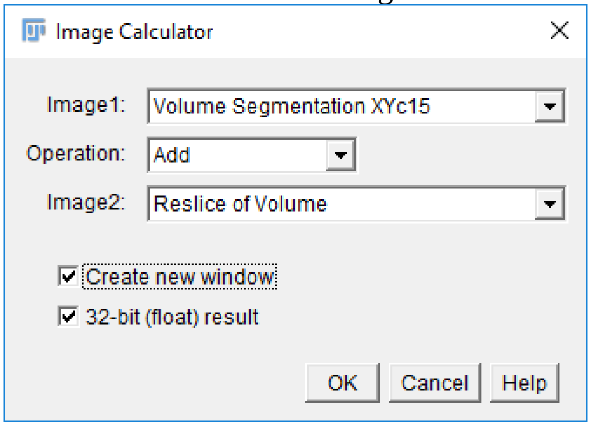


9. Threshold the resulting mask to display the largest pixel values to make sure consensus is maximized. Use the same method as in step 5.
10. Filter the resulting volume from step 9 by 2 pixels in the x,y and z directions using a 3D medium filter with the option Process -> Filters -> Median 3D. This will be the final, resulting segmentation.
11. To visualize the final segmentation in 3D, use the option Plugins -> 3D Viewer.

#### Appendix

Necessary Linux Commands:
ls – lists all files and folders in the current directory.
pwd – gives the full name of the current directory.
cd DIRECTORY – enters the directory specified by the parameter DIRECTORY.
mkdir DIRECTORY – creates a new folder named by the parameter DIRECTORY in the current directory.

**configure.sh**

```
#!/bin/bash
export CLASSPATH=$CLASSPATH:bsh-2.0b4.jar
echo "CLASSPATH: $CLASSPATH"
echo "Make sure bsh-2.0b4.jar is on the end of your CLASSPATH"
module load openjdk/1.8.0_121
java -version
echo "Make sure your java version has 1.8 as the beginning"
```

**generate_swarm_script.sh**

```
#!/bin/bash
function printHelp {
	echo "========================="
	echo "SCRIPT GENERATOR FOR SWARM"
	echo "========================="
	echo "SYNOPSIS"
	echo "./generate_swarm_script.sh [--help] [ INPUT FOLDER ] [ CLASSIFIER MODEL
] [ RESULT MODE ] [ OUTPUT FOLDER ]"
	echo ""
	echo "DESCRIPTION"
	echo "generate_swarm_cript.sh creates the file run.swarm which is the script for the
swarm feature of biowulf"
	echo "Write the parameter options into the script as shown above without the brackets"
	echo "Type --help (with the double dash and no other parameters) to display this
discription message about the script"
	echo "Run the file run.swarm using the command 'swarm -f run.swarm -t 1 -p 2 -g 25'"
	echo ""
	echo "OPTIONS"
	echo "help - if help immediately follows ./main then the description of the program will
be displayed"
	echo "INPUT FOLDER is the name of the folder that contains the input TIFF files"
	echo "CLASSIFIER MODEL - the .model file that Weka uses to run machine learning
classification"
	echo "RESULT MODE - type either 'Probabilities' or 'Segmentation' for the type of
output that you want"
	echo "OUTPUT FOLDER - is the name of the folder that the output will go to"
}
```

```
ARGC=$#
if [ $ARGC -eq 0 ]; then
	echo "ERROR: No parameters present"
	exit 1
fi
if [ $1 == "--help" ]; then
	printHelp
	exit 0
fi
if [ $ARGC -lt 4 ]; then
	echo "ERROR: Not enough parameters present"
	exit 1
fi
if [ $3 != "Probabilities" ] && [ $3 != "Segmentation" ]; then
	echo "ERROR: RESULT parameter incorrectly inputted"
	echo "Your input: $3"
	echo "Please type either Probabilities or Segmentation"
	exit 1
fi
FILES=$1/*.tif
MODEL=$2
RESULT=$3
OUTPUT=$4
CMD="java bsh.Interpreter source.bsh $4 $2 $3"
CMD="./ImageJ-linux64 --ij2 --headless --run source.bsh
'outputDir=\"$4\",modelPath=\"$2\",resultMode=\"$3\""
rm -f run.swarm
touch run.swarm
for f in $FILES; do
	echo "$CMD,inputFile=\"$f\"'" >> run.swarm
done
```

**source.bsh**

```
// @File(label="Output", description="Select the output directory", style="directory") outputDir
// @File(label="Model", description="Select the Weka model to apply") modelPath
// @String(label="Result",choices={"Labels","Probabilities"}) resultMode
// @File(label="Input", description="Select input file", style="file") inputFile

import trainableSegmentation.WekaSegmentation;
import ij.io.FileSaver;
import ij.IJ;
import ij.ImagePlus;
import ij.plugin.Duplicator;
IJ.log( "** Imported libraries. **" );
```

```
// starting time
startTime = System.currentTimeMillis();

// caculate probabilities or segmentation
getProbs = resultMode.equals( "Probabilities" );

// create segmentator
IJ.log( "** Opening Weka segmentation... **" );
segmentator = new WekaSegmentation();
IJ.log( "** Opened Weka segmentation. **" );
// load classifier
IJ.log( "** Importing classifier... **" );
segmentator.loadClassifier( modelPath.getCanonicalPath() );
IJ.log( "** Imported classifier. **" );

// get input image
IJ.log( "** Importing file... **" );
if( inputFile.isFile() )
{
	// try to read file as image
	image = new ImagePlus( inputFile.getCanonicalPath() );
	IJ.log( "** Imported file. **" );
	if( image != null )
	{
		// apply classifier and get results (0 indicates number of threads is auto-detected)
		result = segmentator.applyClassifier( image, 1, getProbs );
		IJ.log( "** Classifier applied. **" );
		// save result as TIFF in output folder
		if ( result != null) {
			slice = new Duplicator().run(result, 1, 1);
			outputFileName = "SEG_" + inputFile.getName().replaceFirst("[.][^.]+$",
"") + ".tif";
			new FileSaver( slice ).saveAsTiff( outputDir.getPath() + File.separator +
outputFileName );
			IJ.log( "** File saved. **" );
		}
		// force garbage collection (important for large images)
		result = null;
		slice = null;
		image = null;
		System.gc();
	}
}
else {
	IJ.log( "Imported address not a file!" );
}
```

```
// print elapsed time
estimatedTime = System.currentTimeMillis() - startTime;
IJ.log( "** Finished processing file in " + estimatedTime + " ms **" );
```
